# Mutations in *pmrB* confer cross-resistance to the LptD inhibitor POL7080 and colistin in *Pseudomonas aeruginosa*

**DOI:** 10.1101/571224

**Authors:** Keith P Romano, Thulasi Warrier, Bradley E Poulsen, Phuong H Nguyen, Alexander R Loftis, Azin Saebi, Bradley L Pentelute, Deborah T Hung

## Abstract

*Pseudomonas aeruginosa* is a major bacterial pathogen for which there is rising antibiotic resistance. We evaluated the resistance mechanisms of *P. aeruginosa* against POL7080, a species-specific, first-in-class antibiotic in phase 3 clinical trials targeting the lipopolysaccharide transport protein LptD. We found resistance mutations in the two-component regulator *pmrB*. Genome-wide transcriptomics and confocal microscopy studies together suggest that POL7080 is vulnerable to the same resistance mechanisms described previously for polymyxins, including colistin, that involve lipid A modifications to mitigate antibiotic cell surface binding.

---

*P. aeruginosa* and other Gram-negative bacteria are becoming increasingly resistant to current antibiotics, and pose a major threat to patients with hospital-acquired infections, compromised immune systems, and chronic pulmonary infections^1-4^. Unfortunately, the discovery of new agents targeting Gram-negative bacteria is especially challenging due to the exclusion of most small molecules by their impermeable cell wall^5^ and array of efflux pumps. Among the last-resort antibiotics currently used to treat severe multi-drug resistant pseudomonal infections are the polymyxin class of cationic antimicrobial peptides (cAMPs), including polymyxin B and colistin (polymyxin E). More recently, the first-in-class antibiotic POL7080, currently in phase 3 clinical trials, was reported with species-specific activity against *P. aeruginosa* by inhibiting the lipopolysaccharide (LPS) transport protein LptD^6-8^. The discovery of POL7080 (and its analogue POL7001) emerged from extensive chemical modifications of the cAMP protegrin-1 (PG-1), in which a beta-hairpin was introduced to create cyclized peptidomimetic analogues^9-10^. The mechanism of action (MOA) of POL7080 and its analogues differ from that of other cAMPs in several key ways. Polymyxins and PG-1 interact with LPS and exhibit broad spectrum anti-microbial activity through self-promoted uptake across the outer membrane, followed by cell lysis through ill-defined mechanisms^11-13^. The LptD inhibitors POL7080 and POL7001, however, have been reported to exhibit a non-lytic MOA through LptD inhibition in *P. aeruginosa* exclusively^6-8^.

To investigate the resistance mechanisms to POL7080 and its analogues, we selected for spontaneously resistant *P. aeruginosa* PA14 mutants by plating mid-log culture on Lysogeny Broth (LB) agar containing 1μM POL7001 (∼15X the liquid MIC). Detailed experimental protocols are outlined in the supplemental materials section. We isolated six independent mutants and confirmed their resistance to POL7080, POL7001, and PG-1 by broth microdilution as described in the Clinical and Laboratory Standards Institute Guidelines^14^. All clones were highly resistant with MIC shifts ranging 5–40 fold relative to PA14 (Table 1), with absolute MICs comparable to four POL7080-resistant clinical isolates reported previously^15^. Whole-genome sequencing revealed 1 to 4 single nucleotide polymorphisms (SNPs) in each mutant relative to wildtype PA14, and all six carried a mutation in a common gene *pmrB* (**Table 1**). PmrB is a histidine kinase and the membrane-bound sensor in the PmrA-PmrB two-component regulatory system in Gram-negative bacteria. In response to low Mg^2+^ levels and periplasmic AMPs, PmrB undergoes conformational changes in its histidine kinase and methyl-accepting (HAMP) domain, leading to autophosphorylation, phosphoryl group transfer to its cognate response regulator PmrA, and downstream activation of transcriptional programs regulating LPS modification^16^. Interestingly, mutations in the PmrA-PmrB have been implicated in polymyxin resistance through upregulation of the lipid A deacylase *pagL* and the *arnBCADTEF-ugd* operon, resulting in LPS modifications that reduce polymyxin binding to the cell surface^17-27^. Notably, one of our isolated mutants, PA14-*pmrB*_*G188S*_, contained an amino acid substitution in the HAMP domain at exactly the same site previously implicated in colistin resistance in *P. aeruginosa* isolates from patients with cystic fibrosis (*pmrB*_*G188D*_)^21^. We thus decided to measure the MICs of colistin against all six resistant mutants and found cross-resistance in all cases with a 4–8 fold shifts in MIC (**Table 1**).

**Table 1:**
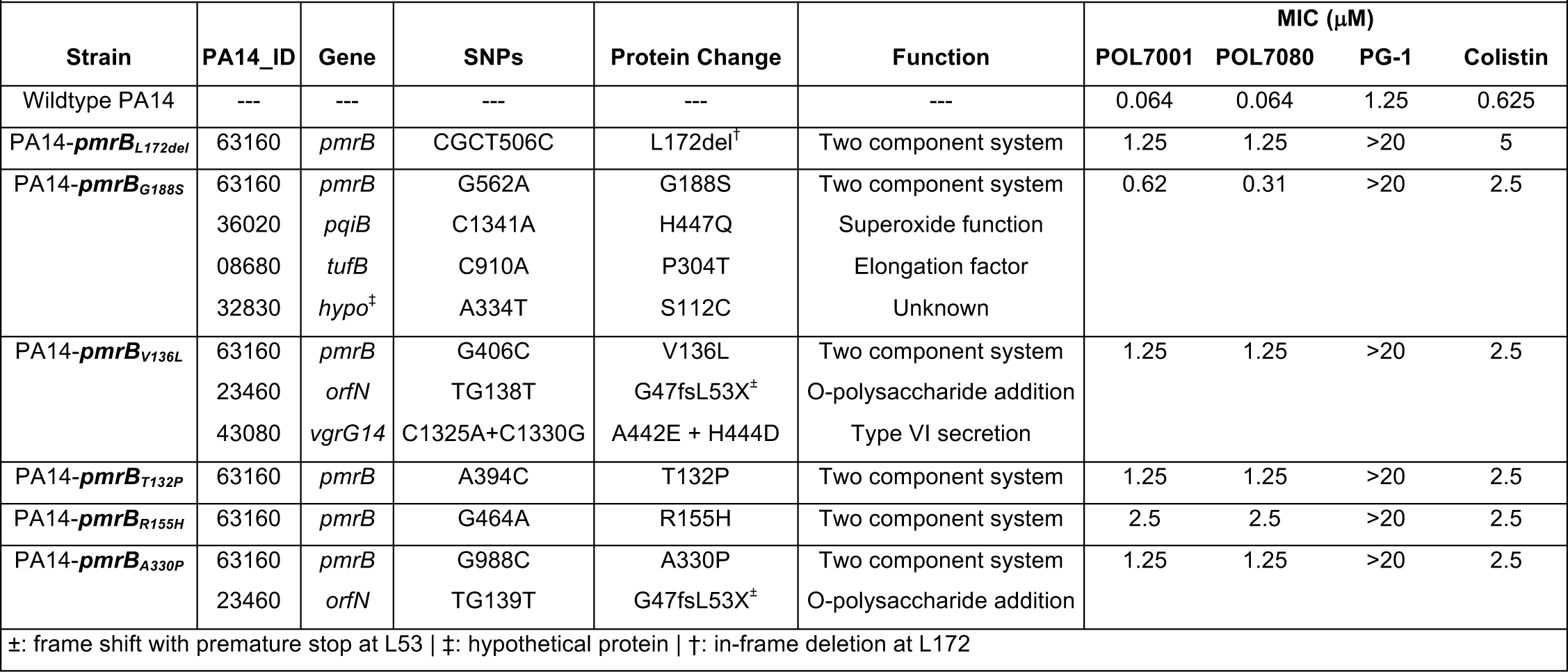
Summary of resistant mutants sequenced after selection with POL7001.

To confirm that alterations in *pmrB* account for the observed resistance to POL7080 and colistin, we introduced a copy of the wildtype allele *pmrB*_*WT*_, and the resistant alleles *pmrB*_*L172del*_ and *pmrB*_*G188S*_, under the control of an arabinose promoter into PA14 at the neutral attTn7 chromosomal site using the pUC18-derived mini-Tn7 integration system^28^. We also introduced the previously reported allele *pmrB*_*G188D*_, conferring colistin resistance^21^, into the wildtype background. MIC assays in the presence of 0.25% (v/v) arabinose demonstrated that all three mutated *pmrB* alleles, but not the wildtype allele, conferred POL7080 and colistin resistance (**Table 2**). Conversely, the introduction of the *pmrB*_*WT*_ allele into the resistant mutant PA14-*pmrB*_*L172del*_ did not restore POL7080 susceptibility with the addition of arabinose, suggesting that resistant *pmrB* alleles were largely dominant over the wildtype allele.

**Table 2:** MICs in 0.25% arabinose after introduction of second *pmrB* alleles into PA14 and *pmrB*_*L172del*_ backgrounds.

|  |  |  |  |
| --- | --- | --- | --- |
| Background strain | attTn7 Allele | POL7080 MIC (μM) | Colistin MIC (μM) |
| PA14 | --- | 0.064 | 0.50 |
| PA14 | *pmrB*<sub>WT</sub> | 0.10 | 0.50 |
| PA14 | *pmrB*<sub>L172del</sub> | 0.60 | >4.0 |
| PA14 | *pmrB*<sub>G188S</sub> | 0.20 | 4.0 |
| PA14 | *pmrB*<sub>G188D</sub> | 0.60 | >4.0 |
| PA14-*pmrB*<sub>L172del</sub> | --- | 1.25 | 5 |
| PA14-*pmrB*<sub>L172del</sub> | *pmrB*<sub>WT</sub> | 0.60 | ND |

We next performed expression analysis (RNA-seq) to investigate the role of *pmrA*-*pmrB* in response to LptD inhibitors. After extracting total RNA from mid-log PA14 treated with 128nM POL7001 (2X the MIC), we prepared RNA-seq libraries using the RNA TagSeq protocol^29^, sequenced samples on an Illumina NextSeq instrument, and analyzed the data using Burrows-Wheeler Aligner^30^ for alignment and DESeq2^31^ to determine differential gene expression. We found that LPS modification genes were significantly upregulated in response to POL7001, including the *pmrA-pmrB*, the lipid A deacylase *pagL*, and entire *arnBCADTEF-ugd* operon responsible for adding 4-amino-4-deoxy-l-arabinose (L-Ara4N) to lipid A (**Fig. 1A**). The aminotransferase *arnB,* catalyzing the final step in L-Ara4N addition, was among the most highly upregulated genes in the entire dataset. We confirmed these findings with quantitative reverse transcription PCR (qRT-PCR) after treating mid-log PA14, or resistant PA14-*pmrB*_*L172de*l_, with either POL7080 or vehicle control (**Fig. 1B**). Relative to control, POL7080 significantly induced *pmrA, arnB* and *pagL* expression in PA14. Notably, *arnB* and *pmrA* transcript levels in untreated PA14-*pmrB*_*L172de*l_ exceeded those in POL7080-treated PA14. Together these data reveal that a signature transcriptional response, the upregulation of key LPS modification genes^17-27, 32^, is constitutively present in the resistant PA14-*pmrB*_*L172de*l_ based on upregulation of key LPS modification genes. These results highlight a common cellular response to LptD inhibitors and polymyxins, and support a shared mechanism by which *pmrB* mutations confer cross-resistance to POL7080 and colistin.

**Figure 1.**
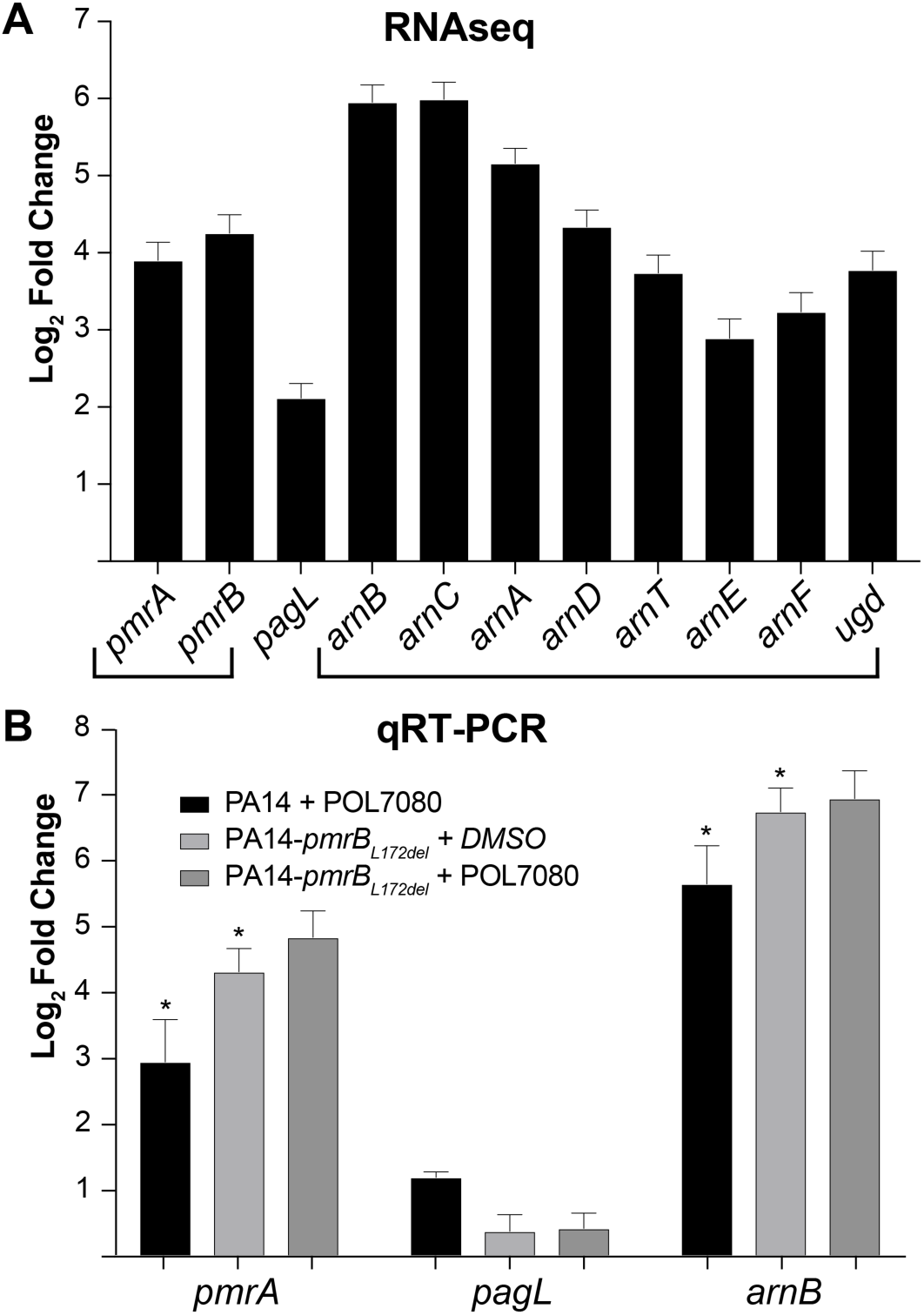
LPS modification genes are upregulated in response to POL7001 and POL7080 treatment and constitutively expressed in the resistant PA14-*pmrB*_*L172del*_ *strain*. (A) RNAseq data shows log_2_ (fold change) in sequencing reads of PA14 after treatment with POL7001 (37°C, 100 minutes) relative to vehicle control. Bracketed genes are located within the same operon. Upregulated genes include the *pmrA*-*pmrB* two-component regulatory genes, lipid A deacylase *pagL*, and the *arnBCADTEF-ugd* operon. (B) After treatment of PA14 and resistant PA14-*pmrB*_*L172del*_ with POL7080 or DMSO control (37°C, 100 minutes), qRT-PCR data shows log_2_ (fold change) in LPS modification gene transcript levels (normalized to *rpoD* expression) relative to PA14 vehicle control. Asterisks indicate paired t-test p-values < 0.03 between POL7080-treated PA14 and PA14-*pmrB*_*L172del*_ vehicle control. In all experiments, error bars represent S.E.M of three biological replicates.

Finally, we investigated whether *pmrB* resistance mutations mitigate POL7080 binding to the cell surface. We synthesized TAMRA-L27-11 (**Fig. S1**), a red-fluorescent analogue of POL7080 with retained inhibitory activity (**Fig. S2**), to probe for differential uptake in PA14-*pmrB*_*L172de*l_ relative to PA14 using confocal microscopy. After treating mid-log PA14 or PA14-*pmrB*_*L172de*l_ with 500nM TAMRA-L27-11, cells were washed, fixed with 4% paraformaldehyde, and stained with 4′,6-diamidino-2-phenylindole (DAPI) for nucleic acid visualization. Red-field and blue-field confocal microscopy showed comparable DAPI staining but over 3-fold reduction in TAMRA-L27-11 uptake in PA14-*pmrB*_*L172de*l_ relative to PA14 (**Fig. 2**), indicating less efficient drug binding at the cell surface of the resistant mutant.

**Figure 2.**
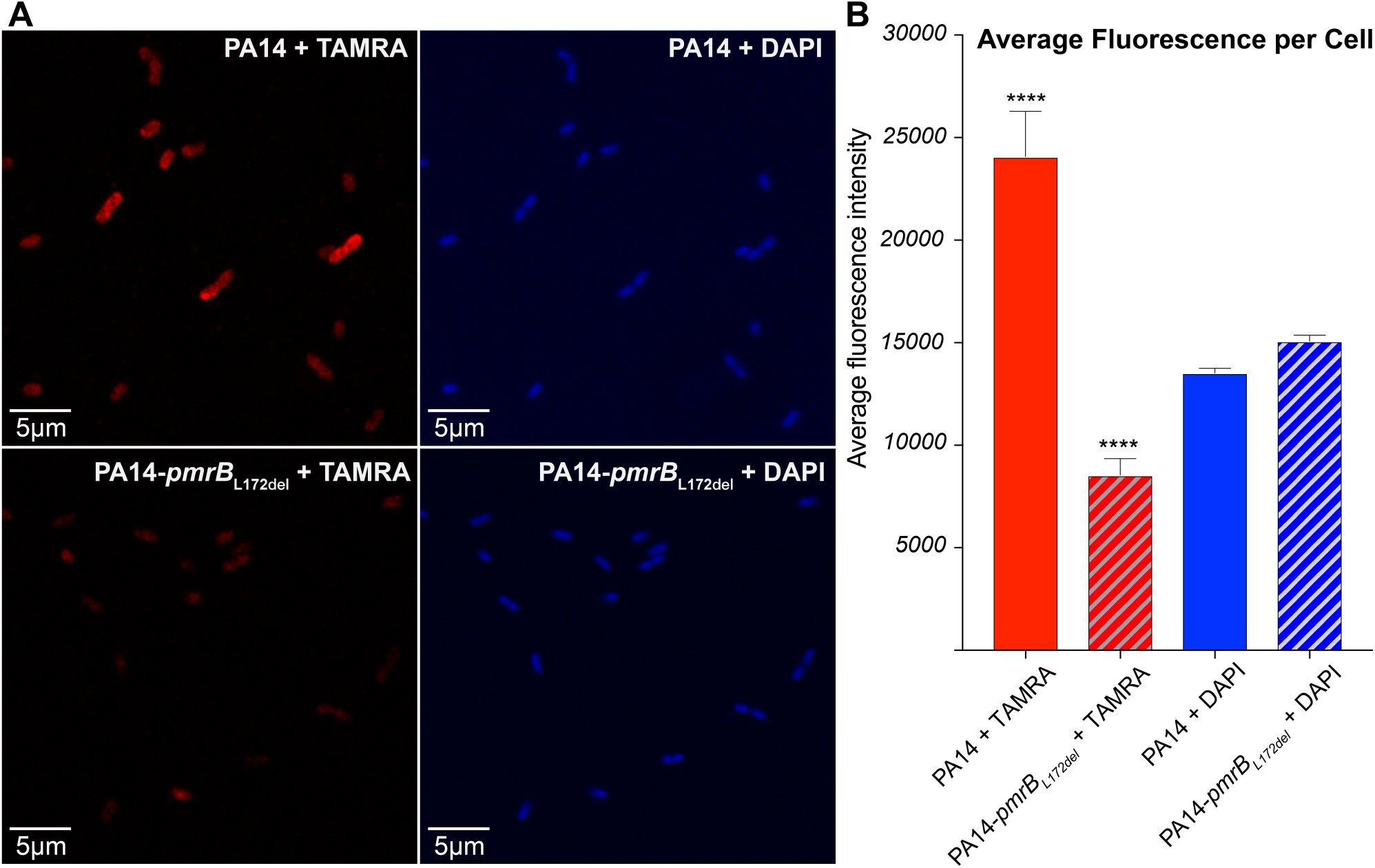
Differential uptake of TAMRA-L27-11 by PA14 versus PA14-pmrB_L172del_. (A) Red-field and blue-field confocal microscopy images of PA14 (top panels) and PA14-pmrB_L172del_ (bottom panels) show reduced TAMRA-L27-11 uptake in PA14-*pmrB*_*L172del*_ relative to PA14. All cells were DAPI stained after treatment with 500nM TAMRA-L27-11 for 120 minutes. (B) Average fluorescence intensities of TAMRA (red bars) and DAPI (blue bars) were calculated for PA14 (solid bars) and resistant PA14-*pmrB*_*L172del*_ (striped bars) using ImageJ software. Error bars represent S.E.M, and asterisks indicate unpaired t-test p-value < 0.0001 between PA14 and PA14-*pmrB*_*L172del*_ after TAMRA-L27-11 treatment.

In summary, we report a series of *pmrB* mutations that confer high-level resistance to POL7080 and moderate cross-resistance to colistin. Expression analysis and confocal microscopy data support a resistance mechanism in which *pmrB* mutations result in LPS modifications by transcriptionally regulating the *arnBCADTEF-ugd* operon, known to result in L-Ara4N addition to LPS, to reduce drug binding to the cell surface. These data align well with known resistance mechanisms to polymyxins, in which LPS modification with L-Ara4N reduces drug binding to the cell surface^20-22^. Our findings suggest that pre-existing colistin resistance may limit the utility of POL7080 in a subset of highly resistant cases of *P. aeruginosa*, and that if successfully developed, POL7080 exposure could inadvertently drive cross-resistance to colistin and other polymyxins.

## Acknowledgements

This work was supported by a generous gift from Anita and Josh Bekenstein and NIH grant 1R01AI117043-04 (D.T.H.). We thank Jonathan Livny, Nirmalya Bandyopadhyay, James Gomez and Roby Bhattacharyya for their helpful discussions. We thank Anilkumar Nair for assistance with confocal microscopy imaging. We declare that we have no conflicts of interest.

