## Supplemental Material for "Mutations in *pmrB* confer cross-resistance to the LptD inhibitor POL7080 and colistin in *Pseudomonas aeruginosa*"

### Supplemental Materials

#### Selecting for POL7001 resistant mutants

PA14 (*Pseudomonas aeruginosa* UCBPP-PA14) cells were grown to mid-log phase ($OD_{600}$ 0.4–0.6) in Lysogeny Broth (LB). Cells were pelleted at room temperature with centrifugation (5000g, 10 minutes) and resuspended in LB to yield $1 \times 10^9$ CFU/mL ($OD_{600}$ 1.0). $\sim 10^8$ CFU were plated onto agar plates containing 1μM POL7001 (~15X its MIC in liquid culture). Six resistant mutants were isolated after incubation at 37°C for 16 hours, which were further grown in 5mL LB with 200nM POL7001. Total DNA was extracted using the DNeasy Blood and Tissue Kit (Qiagen, Hilden Germany), and subsequently quantified using the High Sensitivity Quant-iT dsDNA Assay Kit (Thermo Fisher Scientific, Waltham MA). Indexed paired-end libraries were generated using the Nextera XT DNA library preparation kit (Illumina, San Diego CA). 25μL of index DNA libraries were mixed with 15μL Agencourt AMPure XP beads (Beckman Coulter, Pasadena CA) in a 0.5mL deep well block (Thermo Fisher Scientific, Waltham MA), and after equilibrating on a 96S Super Magnet (Alpaqua Engineering, Beverly MA) for 2 minutes, the supernatants were removed. Samples were twice washed with 200μL of 80% (v/v) ethanol, and after drying on the magnet stand for 15 minutes, purified DNA was eluted with 25μL water. DNA quality and quantity were determined using the D5000 ScreenTape with the 2200 TapeStation (Agilent, Santa Clara CA). Samples were diluted to a final concentration of 6.8ng/μL, and pooled for sequencing on the MiSeq instrument (Illumina, San Diego CA). Genomes and annotations were obtained from www.pseudomonas.com[1], and Illumina reads were mapped to the PA14 genome and single nucleotide polymorphisms were identified using the Pithon program suite previously reported[2].

#### MIC determination of POL7001-resistant mutants

MIC experiments were conducted in biological triplicate in 384-well microplates (Nunc 384-well clear polystyrene plates, Thermo Fisher Scientific, Waltham MA) using the standard microdilution broth method adapted from the Clinical and Laboratory Standards Institute Guidelines[3]. For colistin, 2-fold serial dilutions were performed in 1% (v/v) DMSO in LB at 2X the desired assay concentrations. For POL7001, POL7080 and PG-1, 2-fold serial dilutions were performed in dimethyl sulfoxide (DMSO) at 200X the desired assay concentrations, after which samples were diluted 1:100 into LB. 30μL of serially diluted drug solutions were mixed with equal volume mid-log bacterial culture to yield final conditions of $5 \times 10^5$ CFU/mL bacteria with drug in LB with 0.5% (v/v) DMSO. Microplates were incubated in a humidity chamber for 16 hours at 37°C, after which time the $OD_{600}$ was measured in the Spark Multimode Reader (Tecan, Männedorf Switzerland). Data were plotted and analyzed using GraphPad Prism8 software to determine the MICs, defined as the minimal drug concentrations required for complete inhibition of bacterial growth.

**Introduction of *pmrB* alleles at the attTn7 chromosomal site**

A copies of the wildtype *pmrB* allele, POL7080-resistant *pmrB* alleles L172del and G188S, and the colistin-resistant *pmrB* allele G188D[4] were introduced into PA14 at the neutral, naturally-evolved attTn7 chromosomal site using the mini-Tn7 system described previously in *P. aeruginosa*[5]. When possible, the desired *pmrB* alleles were amplified from the genomic DNA of its corresponding mutant and inserted into the pUC18-derrived mini-Tn7 integration vector (with gentamicin cassette and AraC-*araBAD* promoter system) using standard PCR and Gibson Assembly protocols (New England Biolabs, Ipswich MA)[6]. Because no mutants were available with the *pmrB* allele G188D, the substitution was instead engineered into the wildtype *pmrB* mini-Tn7 integration vector using the Q5 Site-Directed Mutagenesis Kit (New England Biolabs, Ipswich MA). Ultimately, each mini-Tn7 integration vector constructed contained the desired *pmrB* allele (wildtype, L172del, G188S, or G188D) under control of the AraC-*araBAD* promoter[7], which encodes the *araC* repressor allowing for titratable *pmrB* gene expression in response to arabinose. After transformation into 10-beta *Escherichia coli* (New England Biolabs, Ipswich MA), bacterial conjugation was performed on cellulose membranes by spotting 20μL of a 1:2:2:2 mixture of the recipient strain (PA14 or PA14-$pmrB_{172del}$), 10-beta cells harboring the desired *pmrB* integration vector, and helper strains pRK2013 and pTNS3 (encoding the machinery necessary for pseudomonal plasmid uptake and subsequent integration at the attTn7 site). After mating at 37°C for 10 hours, cells were resuspended in 400μL LB and selected on LB agar containing 30μg/mL gentamicin and 15μg/mL irgasan, permitting growth of only PA14 cells containing the *pmrB* gene (and gentamicin cassette) integration. Colonies were isolated and *pmrB* gene inserts were confirmed by fragment sizing after colony PCR. PCR products of the appropriate size were also recovered using the Gel Extraction Kit (Qiagen, Hilden Germany), inserted into Zero Blunt TOPO vector (Thermo Fischer Scientific, Waltham MA), and confirmed by whole plasmid sequencing.

**MIC determination of PA14 strains with second arabinose-inducible *pmrB* alleles**

MIC experiments were conducted in biological triplicates in 384-well microplates (Nunc 384-well clear polystyrene plates, Thermo Fisher Scientific, Waltham MA) as detailed above using PA14 clones containing the second alleles $pmrB_{WT}$, $pmrB_{L172del}$, $pmrB_{G188S}$, and $pmrB_{G188D}$. MIC assays were also performed using PA14-$pmrB_{L172del}$ containing a second $pmrB_{WT}$ allele. Starter cultures were diluted into LB with 0.25% (v/v) arabinose. For colistin, 2-fold serial dilutions were performed in LB with 0.5% (v/v) DMSO and 0.25% (v/v) arabinose at 2X the desired assay concentrations. For POL7080, 2-fold serial dilutions were performed in DMSO at 400X the desired assay concentrations, after which samples were diluted 1:200 into LB with 0.25% (v/v) arabinose. For each arabinose condition, 30μL of mid-log cultures were mixed with equal volume of POL7080 (or colistin) to yield a final condition of $5x10^5$ CFU/mL bacteria with drug in 0.25% (v/v) DMSO in LB with 0.25% (v/v arabinose). Microplates were incubated in a humidity chamber for 16 hours at 37°C, after which time the $OD_{600}$ was measured in the Spark Multimode Reader (Tecan, Männedorf Switzerland). Data were plotted and analyzed using GraphPad Prism8 software to determine the MICs.

**RNAseq experiments**

RNAseq experiments were carried out in biological triplicate in 384-well microplates (Nunc 384-well clear polystyrene plates, Thermo Fisher Scientific, Waltham MA). POL7001 was dissolved in DMSO at 200X the working concentration, and then diluted 1:100 into LB, yielding 2X drug solution in 1% (v/v) DMSO in LB. PA14 cultures were grown with shaking (37°C, 250rpm) to mid-log phase in LB. 30μL of mid-log cultures were mixed with equal volume of 2X POL7001 solution, yielding final conditions containing of $2x10^8$ CFU/mL PA14 with 128nM POL7001 in 0.5% (v/v) DMSO in LB. Cells were treated at 37°C for 100 minutes without shaking in a humidity chamber. Samples were then mixed with 30μL of 3X RNAgem Blue Buffer (Zygem, Charlottesville VA), and chemically lysed by incubation in a thermocycler at 75°C for 10 minutes. Total RNA was then extracted using the Direct-zol kit (Zymo Research, Irvin CA), and RNA quality and quantity were analyzed using the RNA ScreenTape with the 2200 TapeStation (Agilent, Santa Clara CA). RNA-seq libraries were prepared using the RNA TagSeq protocol previously described[8], and samples were sequenced samples on a NextSeq instrument (Illumina, San Diego CA). Transcriptional data were analyzed using the Burrows-Wheeler Aligner[9] for alignment and DESeq2[10] to determine genes differentially expressed.

**qRT-PCR experiments**

qRT-PCR experiments were carried out in biological triplicate in a Nunc 96-well 2mL Deep Well Block (Thermo Fisher Scientific, Waltham MA). POL7080 was dissolved in DMSO at 200X the working concentration, and then diluted 1:100 into LB, yielding 2X drug solution in 1% (v/v) DMSO in LB. PA14 and resistant PA14-*pmrB*$_{L172del}$ cultures were grown with shaking (37°C, 250rpm) to mid-log phase in LB. 125μL of mid-log cultures were mixed with equal volume of either 2X POL7080 solution or 1% DMSO vehicle control, yielding final conditions containing of $2x10^8$ CFU/mL bacteria with 100nM (or 0nM) POL7080 in 0.5% (v/v) DMSO in LB. Cells were treated at 37°C for 100 minutes without shaking in a humidity chamber. Samples were then pellets by centrifugation (5,000g, 10 minutes), and 150μL of supernatant was discarded. After resuspension in the remaining supernatant, 90μL of culture were transferred to an Axygen 96-well PCR plate (Corning Incorporated, Corning NY), containing 10μL of 10X RNAgem Blue Buffer (Zygem, Charlottesville VA). Chemically lysis was performed by incubation in a thermocycler at 75°C for 10 minutes. Total RNA was then extracted using the Direct-zol kit (Zymo Research, Irvin CA), and RNA quality and quantity were analyzed using the RNA ScreenTape with the 2200 TapeStation (Agilent, Santa Clara CA). Complementary DNA (cDNA) libraries were generated using the qScript cDNA synthesis kit (QuantaBio, Beverly MA). All qRT-PCR reactions were carried with the ViiA 7 Real-Time PCR system (Applied Biosystems, Foster City CA). Primer pairs were designed for *pmrA*, *arnB*, *pagL* and *rpoD* genes and purchased from Integrated DNA Technologies (**Table S1**). cDNA, primer pairs, and iTaq Universal SYBR Green Supermix (Bio-Rad Laboratories) were mixed according to the iTaq Universal SYBR Green Supermix protocol. 10μL of PCR reaction mixtures were transferred to 384-well thin-wall hard-shell PCR plates (Bio-Rad Laboratories, Hercules CA). Cycle conditions were based on the iTaq SYBR Green mix protocol for the BIO-RAD CFX384 system as follows: polymerase activation and DNA denaturation at 95ºC for 20 seconds for cDNA, or 5 minutes for genomic DNA; 40 cycles for amplification (denaturation at 95ºC for 5 seconds, followed by annealing at 63ºC for 30 seconds, followed by extension at 72ºC for 2 minutes). A series of seven 10-fold dilutions of PA14 control genomic DNA (initial stock at 40ng/μL) were performed, and standard curves were generated to determine the conversion factor of DNA copies per PCR reaction, assuming 1ng of PA14 genomic DNA equaled $1.4x10^5$ DNA copies. For analysis, gene transcript levels were normalized to those of the housekeeping *rpoD* gene, and subsequently normalized to untreated PA14 transcript levels to determine fold-change in gene expression relative to untreated PA14.

**Table S1: Optimal qRT-PCR primer pairs for genes *pmrA*, *arnB*, *pagL*, and *rpoD*.**

| Gene | Forward Primer | Reverse Primer |
|---|---|---|
| ***pmrA*** | AAGGCGATACCGTGGAATG | CAGGTTGCGCAGGATGT |
| ***arnB*** | CTGGCACCTGTTCATCCTG | AGGTGACTGGCGATGAAATG |
| ***pagL*** | GATGCGGGCTACACCTATTG | GCCTCGATGAATGGCTTGAT |
| ***rpoD*** | GATTTCCATCGCCAAGAAGT | CACGACGGTATTCGAACTTG |

**Synthesis of TAMRA-modified cyclic peptide L27-11 (TAMRA-L27-11)**

*Synthesis of parent peptide*

The parent cyclic peptide L27-11 was synthesized mostly as previously described[11]. The uncyclized linear peptide was prepared by a combination of batch synthesis and manual flow peptide synthesis ($N^{\alpha}$-Fmoc-protected amino acids from Chem-Impex International)[12]. Except where specified, all reagents in this procedure were purchased from Sigma-Aldrich. In batch, Fmoc-L-Pro-OH (1mmol) was coupled to 150 mg 2-chlorotrityl chloride polystyrene resin (200–400 mesh, loading: 1.14mmol/g, Chem-Impex International) in the presence of diisopropylethylamine (DIEA, 0.3mL) in dichloromethane (DCM, 2.5mL). Manual flow peptide synthesis followed the standard 3-minute cycle at 60°C previously described. Initial deprotection of $N^{\alpha}$-Fmoc was performed with 6.6mL 20% (v/v) piperidine in N,N-dimethylformamide (DMF) delivered at 20mL/min over 20 seconds. Subsequent coupling was performed by delivering 1mmol Fmoc-Xaa dissolved in 2.5mL of 0.4M 2-(7-Aza-1H-benzotriazole-1-yl)-1,1,3,3-tetramethyluronium hexafluorophosphate (HATU, Chem-Impex International) in DMF and 0.5mL DIEA. Between coupling and deprotection steps, resin was washed for 1 minute by delivering 20mL of DMF at 20mL/min. Final amino acid coupling was performed in batch. Fmoc-$N^{\varepsilon}$-azide-L-Lysine-OH (1mmol) was dissolved in 2.5 mL of 0.4M HATU in DMF and 0.5mL DIEA, and added to resin. Subsequent Fmoc deprotection was performed by addition of 20% (v/v) piperidine in DMF (5mL). Cleavage from resin was performed via addition of 1% (v/v) TFA in DCM for 5 minutes. The eluate was transferred and neutralized with 1mL DIEA. Cleavage and neutralization was performed twice. Solvent was then removed under high vacuum. To cyclize, the resulting side-chain protected linear peptide was resuspended in DMF (85mL), and HATU (1mmol, 6 eq.), 1-hydroxy-7-azabenzotriazole (HOAt, 1mmol, 6 eq., Chem-Impex International), and 2,4,6-collidine (2.6mmol, 15 eq.) were added. The reaction was stirred for 72 hours, and DMF was then removed under high vacuum. Crude cyclic peptide was resuspended in DCM (20mL) and washed with 10% (v/v) acetonitrile (MeCN) in water containing 0.1% trifluoroacetic acid (TFA) twice. DCM fraction was transferred to a new vessel and DCM was removed under high vacuum. Crude cyclic peptide was then subjected to global deprotection via resuspension in TFA/thioanisole/water/phenol/1,2-ethanedithiol (82.5:5:5:5:2.5 v/v, 5mL) and incubation at room temperature for 2 hours. Deprotected peptide was precipitated from cleavage solution via trituration with ice cold diisopropyl ether. Crude cyclic peptide was washed twice with ice cold diisopropyl ether, dried, and conjugated to tetramethyl rhodamine (TAMRA).

*Fluorophore conjugation and purification of TAMRA-L27-11*

Crude parent peptide was conjugated to TAMRA using strain-promoted copper-free azide-alkyne click chemistry. Azide-containing parent peptide L27-11 (3mg) was resuspended in 1X phosphate buffered saline (PBS), and dibenzocyclooctyne-PEG4-TAMRA (3 mg, 2 eq., Sigma-Aldrich) resuspended in 5% (v/v) DMSO in 1X PBS was added to peptide, and mixture was incubated at room temperature for 2 hours. Reaction was then purified by preparative RP-HPLC (Agilent Zorbax SB C3 column: 9.4 x 250mm, 5μm) with a gradient of 5-55% MeCN in water with 0.1% TFA, run at 0.5% B/min. Pure fractions were pooled and lyophilized, and final purity was confirmed by LC-MS (**Figure S1**).

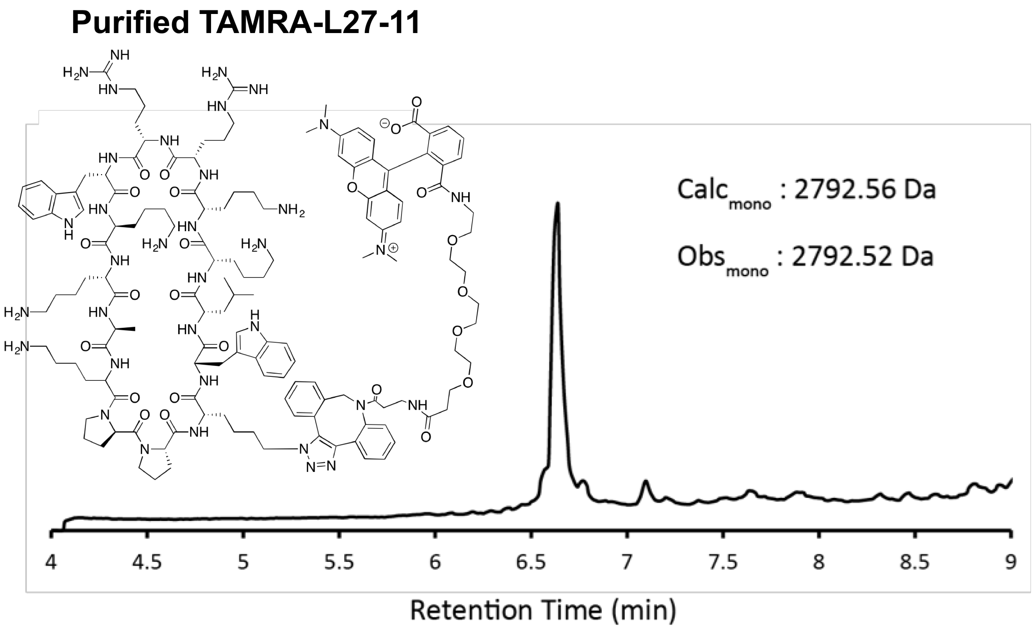


**Figure S1. Chemical structure and LC-MS analysis of RP-HPLC purified TAMRA-modified cyclic peptide L27-11.** Calculated and observed masses are monoisotopic, with total ion chromatogram shown.

**TAMRA-L27-11 Growth Curves**

Growth curves in the presence of TAMRA-L27-11 were conducted in biological triplicate in 384-well microplates (Nunc 384-well clear polystyrene plates, Thermo Fisher Scientific, Waltham MA). TAMRA-L27-11 was dissolved in DMSO at 200X the working concentration, and then diluted 1:100 into LB, yielding 2X TAMRA-L27-11 solution in LB with 1% (v/v) DMSO. PA14 and PA14-*pmrB*$_{L172del}$ cultures were grown with shaking (37°C, 250rpm) to mid-log phase in LB. 30μL of mid-log bacterial cultures were mixed with equal volume of 2X TAMRA-L27-11 solution to yield a final bacterial concentration of $5 \times 10^5$ CFU/mL in 20μM TAMRA-L27-11 in LB with 0.5% (v/v) DMSO. Microplates were incubated in a large humidity cassette at 37°C and the $OD_{600}$ was measured every 30 minutes using the microplate reader (Spark multimode microplate reader, Tecan). Data were plotted and analyzed using GraphPad Prism8 software to generate growth kinetic curves (**Figure S2**).

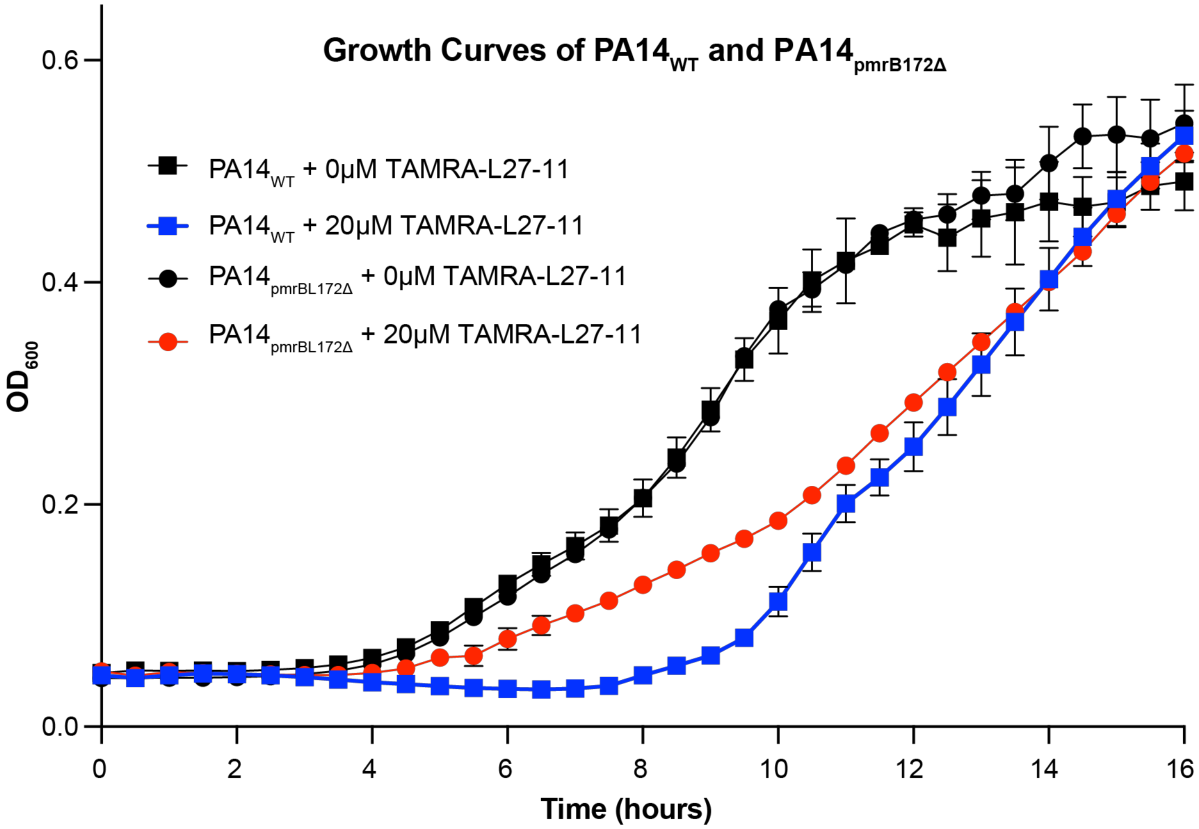


**Figure S2. Differential growth kinetics of PA14-*pmrB*$_{L172del}$ in the presence of TAMRA-L27-11.** PA14-*pmrB*$_{L172del}$ demonstrates a less pronounced growth delay than PA14 in the presence of 20μM TAMRA-L27-11.

**Confocal Microscopy**

PA14 and PA14-$pmrB_{L172del}$ cultures were grown with shaking (37°C, 250rpm) to mid-log phase in LB. TAMRA-L27-11 was dissolved in DMSO at 200X the working concentration, and then diluted 1:100 into LB, yielding 2X TAMRA-L27-11 solution in LB with 1% (v/v) DMSO. 500μL of mid-log bacteria were mixed in culture tubes with equal volume of TAMRA-L27-11 to yield a final bacterial concentration of $1x10^8$ CFU/mL in 500nM TAMRA-L27-11 in LB with 0.5% (v/v) DMSO. After incubation with shaking (37°C, 250rpm) in the dark for 120 minutes, cells were pelleted and twice washed with PBS. All wash steps were carried out at room temperature by resuspension in 500μL PBS, centrifugation (4000g, 10 minutes), and careful removal of the supernatant without pellet disruption. Cells were next fixed by treatment with 4% (v/v) paraformaldehyde in PBS in the dark for 30 minutes on ice. Cells were washed once with PBS, resuspended in 100μL of 100μM 4′,6-diamidino-2-phenylindole (DAPI) dissolved in PBS, and incubated at room temperature for 20 minutes in the dark. Cells were again washed and resuspended in 20μL PBS. On clean microscope slides, 1μL of fixed cells were mixed with 5μL of ProLong Gold Antifade Reagent (Invitrogen, Carlsbad CA). Cover slips were applied and the samples were allowed to dry in the dark at room temperature for at least 24 hours. Blue and red-field confocal microscopy images were collected using the Zeiss Confocal Microscope. For analysis, the average intensities of 20–30 cells were quantified using ImageJ software to calculate mean DAPI and TAMRA fluorescence intensities of PA14 and PA14-$pmrB_{L172del}$ after TAMRA-L27-11 treatment.
